# Sensitive periods for white matter plasticity and reading intervention

**DOI:** 10.1101/346759

**Authors:** Jason D. Yeatman, Elizabeth Huber

## Abstract

As a child matures, some brain circuits stabilize while others remain plastic. However, the literature on maturational changes in the brain’s capacity for experience-dependent plasticity is primarily based on experiments in animals that mature over dramatically different time-scales than humans. Moreover, while principles of plasticity for sensory and motor systems might be conserved across species, the myriad of late-developing and uniquely human cognitive functions such as literacy cannot be studied with animal models. Here we use an intensive reading intervention program, in combination with longitudinal diffusion MRI measurements in school-aged children with dyslexia, to investigate the sensitive period for white matter plasticity and literacy learning. We find that the intervention induces large-scale changes in white matter diffusion properties, and improvements in reading scores, but that the magnitude and time-course of plasticity does not depend on the subject’s age. Thus, we conclude that, for the intensive, one-on-one reading intervention program employed here, if a sensitive period exists, it does not end before middle school.

## Introduction

Younger brains are more plastic. That the brain’s capacity for plasticity diminishes with age is commonly held as an axiom of development and carries important implications for education and the treatment of developmental disorders. For example, developmental dyslexia is believed to be rooted in neuroanatomical differences within well-characterized brain circuits (Langer et al., 2015; Ozernov-Palchik and Gaab, 2016; Saygin et al., 2013; Vandermosten et al., 2015; Wandell and Yeatman, 2013; Yeatman et al., 2012a) and interventions intended to remediate these differences are presumed to be more effective in younger, compared to older, children (Gabrieli, 2009; Gabrieli et al., 2015).

The theory that plasticity diminishes over development takes different forms, ranging from strict critical periods, which define windows of development during which specific experiences can shape the structure of a neural circuit (Hubel et al., 1977; Wiesel and Hubel, 1965, 1963), to sensitive periods, in which circuits exhibit varying degrees of plasticity and propensity for experience dependent change (Blakemore, 1988; Meltzoff et al., 2009). Many aspects of cognition may be subject to broad sensitive periods (e.g., pre-puberty), during which large swaths of cortex remain malleable to environmental demands until rising hormone levels stabilize the relevant circuits (Piekarski et al., 2017a) (reviewed in (Piekarski et al., 2017b)). A wealth of research in model organisms, including mice and non-human primates, has identified cellular changes (e.g., the refinement of ocular dominance columns in primary visual cortex (Hubel et al., 1977; Katz and Shatz, 1996; Tagawa et al., 2005; Wandell and Smirnakis, 2009)) that only occur during isolated developmental windows, and mechanisms (e.g., perineuronal nets (Hensch, 2005; Werker and Hensch, 2014)) that govern the transition from plasticity to stability for a circuit. A wealth of research in humans has identified aspects of learning (e.g., the sound inventory of language (Kuhl, 2004; Meltzoff et al., 2009)) that become more difficult with age. However, to our knowledge, there is no data directly linking changes in the human capacity for learning high-level cognitive functions to changes in the brain’s capacity for experience-dependent structural plasticity. While it is appealing to assume that the timing of critical/sensitive periods discovered in animal models might generalize to human, there are dramatic differences in the time-course of maturation across species. For example the myelination process is prolonged by more than a decade in humans compared to other primate species, even after adjusting for developmental milestones such as puberty (Miller et al., 2012). Therefore, plasticity in humans, particularly for the white matter, might not be subject to the same constraints as other species, highlighting the importance of understanding the principles governing plasticity in the human brain.

Two principal challenges to studying experience-dependent plasticity in humans are, first, establishing an experimental paradigm that is appropriate for human research subjects and capable of inducing large-scale structural changes in the brain and, second, developing non-invasive measurements that are sensitive to changes in cellular properties of human brain tissue. Our previous work demonstrated that combining an intensive reading intervention program with longitudinal diffusion-weighted magnetic resonance imaging (dMRI) measurements in children with dyslexia is a powerful paradigm for studying experience-dependent changes in the white matter (Huber et al., 2018). In a sample of 24 children between 7 and 13 years of age, eight weeks of targeting training in reading skills caused large-scale changes in tissue properties for multiple anatomical tracts (Cohen’s *d* = 0.5-1.0 across different white matter tracts), that were coupled to large improvements in reading skills (Cohen’s *d* = 0.5-1.0 across different reading tests). Here we capitalize on this paradigm, and a larger sample of subjects (N=34), to test the hypothesis that there is a sensitive period for this circuit and that the amount of intervention-driven plasticity measured in the white matter depends on the age of the subject. Specifically, we consider three hypotheses:

1. Younger subjects will show larger intervention-driven changes in diffusion properties compared to older subjects. The magnitude of change for each subject will be computed as (1) the difference between the pre- and post-intervention measurement sessions and (2) a linear fit summarizing the rate of change in diffusion properties as a function of intervention hours.
2. Younger subjects will show more rapid changes compared to older subjects, irrespective of the absolute magnitude of change. The timescale of change for each subject will be computed as (1) the growth rate observed between the first 2 sessions and (2) the percentage of overall change that occurs within the first three weeks of the intervention.
3. Alternatively, we may observe the same magnitude and time-course of experience-dependent white matter plasticity over this age range (7-13 years of age). Such a finding would imply that, for the training paradigm and age range studied here, age is not a major factor in determining a child’s response to intervention, and that the sensitive period for these white matter networks extends from early elementary-school into young adulthood.

## Methods

### Pre-registration

Understanding maturational changes in plasticity that occur over the course of elementary school is an important scientific challenge with practical implications for education practice. Our previous work established an experimental paradigm (intensive, one-on-one reading intervention program) and measurement protocol (longitudinal diffusion MRI measurements) for quantifying experience-dependent plasticity in human white matter (Huber et al., 2018). A power analysis based on these data confirmed that we have the statistical power to detect meaningful maturational differences in experience-dependent plasticity, if such differences exist (see **Figure 1**). However, if the data indicate that older (middle-school aged) children show the same large-scale changes in white matter as younger (1st and 2nd grade) children, it would revise our preconceived notions about brain maturation, plasticity, and learning in an academic setting. Therefore, the outcome of the proposed investigation will answer fundamental scientific questions irrespective of the results.

**Figure 1:**
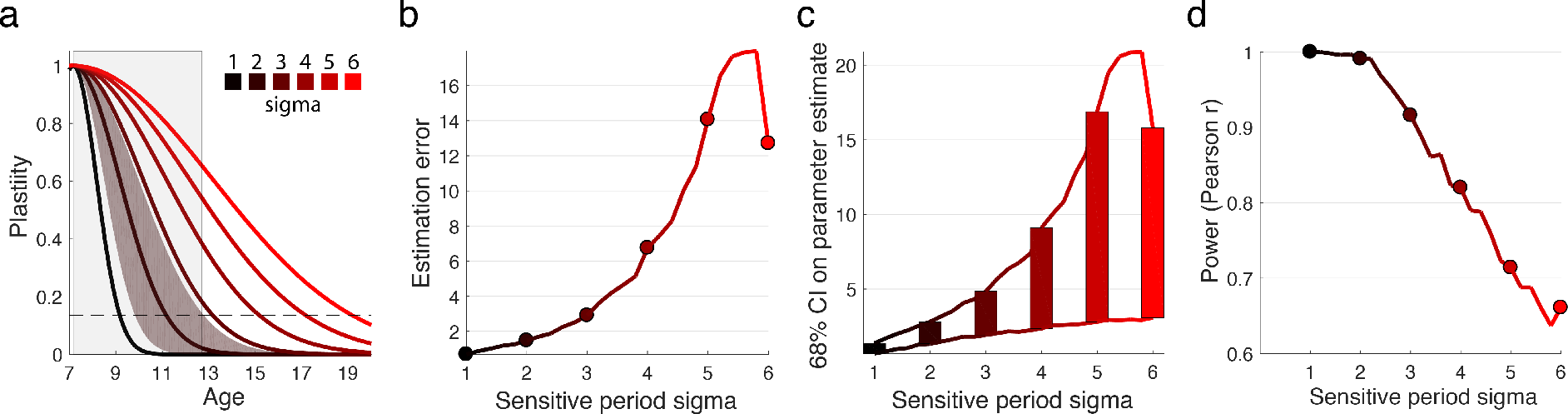
Simulation of sensitive periods. (a) Sensitive periods are modeled as a Guassian function where the width of the Gaussian (sigma) defines the window of plasticity (see simulateAgeEffect.m). The light gray box denotes the age range of subjects in our study. Simulated data was generated from the Gaussian model by adding noise, where the amount of noise was calibrated based on the session-to-session variability of the measurements in the absence of intervention (estimated from data in control subjects). Six curves are illustrated for potential models ranging from sigma=1 to sigma=6 and the standard error of the model fit to the simulated data is illustrated for one curve (sigma=2). (b) The average error on the sigma parameter estimate is shown for different potential models given the signal to noise ratio of our data. The continuous line shows increments of 2.4 months. (c) Confidence intervals are displayed based on models with different sigma parameters. (d) Based on simulating data from different sensitive period models, we calculate the probability of detecting a significant correlation (p < 0.05) between plasticity and age. Simulation code is available at: https://github.com/yeatmanlab/plasticity

We believe that pre-registration is important since our conclusions can depend on how we operationalize our hypotheses and process our data. Hypothesizing after knowing the results, or “HARKing”, has recently been highlighted as a major issue contributing to a publication bias in the biomedical and social sciences (Munafò et al., 2017). Munafo and colleagues maintain that practices such as HARKing are often subconscious and unintentional, reflecting well-known psychological biases that are difficult to avoid. Hence, in line with their perspective, we acknowledge the possibility that our approach to data analysis could change in a manner that is more likely to confirm a statistical relationship between age and plasticity, rather than a lack thereof. While registration prior to data collection is optimal, there are many cases (such as the present one) where it is not feasible. In our case, studying maturational changes in the brain’s capacity for plasticity first required establishing the sensitivity of our measurement protocol, and determining the pattern of changes that are induced by the intervention. Our previous work optimized our analysis pipeline to reliably measure white matter plasticity in an individual and made pre-registration possible for the present study. Thus, the data acquisition and analysis methods are identical to those reported in (Huber et al., 2018) with the addition of 10 new intervention subjects that were run as a replication cohort.

Our methodology for pre-registration was as follows. The complete introduction and methods section were written, and posted as a preprint on bioRxiv, before analyzing the data (June 13, 2018; https://www.biorxiv.org/content/early/2018/06/13/346759). The methodological details and statistical comparisons were designed based on our previous work (Huber et al., 2018), and the hypotheses were defined at the end of the introduction to avoid HARKING. The only changes to the introduction and methods between the pre-registration and the final version were: (1) new title, (2) minor edits to improve the flow of the paper, (3) clarifications of details based on feedback we received on the preprint, (4) the addition of the present paragraph describing our pre-registration methodology. To determine whether we had sufficient statistical power to detect age-dependent changes in plasticity we posited a model of sensitive periods and ran a simulation based on the data released in (Huber et al., 2018) (**Figure 1**; data and simulation code available at: https://github.com/yeatmanlab/plasticity). Since the current pre-registration policies at most journals require registration before data collection starts, we were unable to formally pre-register this study with a journal. Thus, in order to obtain reviews of our methodology and reasoning before analyzing the age-dependence of our previously reported effects, we: (1) sent the link to the preprint to knowledgeable experts in the field to solicit feedback, and (2) posted the link on social media to solicit anonymous reviews from the general public. We then revised the manuscript, completed all of our pre-defined analyses, and pursued additional post-hoc analyses where the results warranted additional exploration. Post-hoc analyses are noted as such in the Results section.

### Modeling sensitive periods and determining statistical power for planned analyses

A sensitive (or critical) period is a window of development during which a circuit is particularly responsive to environmental inputs (Werker and Hensch, 2014). As the balance between excitatory and inhibitory signaling changes, the capacity for plasticity within a circuit increases, marking the onset of a sensitive period. The sensitive period remains open until molecular brakes close the window for plasticity. However, this developmental period need not have a strict boundary, and may instead gradually close, leading to diminished plasticity over a period of months or years. We therefore model this time-course as a Gaussian function where the mean of the Guassian (μ) marks the age (x) of peak plasticity, the height of the Gaussian (β) denotes the magnitude of plasticity that is possible, and the width of the Gaussian (σ) is used to estimate the window of development during which the sensitive period is open.

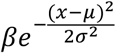

If we assume that the sensitive period for literacy is near its peak at the beginning of elementary school when children begin formal reading instruction, then a narrow sensitive period (e.g., σ=1, black curve in **Figure 1a**) would imply that the window for plasticity in the reading circuitry is shut by 3^rd^ or 4^th^ grade, whereas a broad sensitive period (e.g., σ=6, red curve in **Figure 1a**) would imply that the circuit remains plastic through young adulthood. Based on these hypothetical sensitive periods, ranging from σ=1 to σ=6, we simulate our precision for estimating model parameters (**Figure 1b,c**) and detecting a significant correlation between plasticity and age (**Figure 1d**, based on effects reported in ref (Huber et al., 2018)).

Huber et al., (2018) reported the amount of change measured in the white matter over the course of a tightly controlled and intensive reading intervention program delivered to elementary school children with dyslexia. If we operationalize white matter plasticity as the magnitude of change in mean diffusivity (MD) measured over the course of the intervention program, then we can scale each curve such that the mean value over the measured age range (gray shading **Figure 1a**) is equal to the average MD change reported in the sample. The standard deviation of MD change measured in the control group (which did not undergo the intervention and did not, on average, show any change over the measurement period) is used as an estimate of noise. We then: (1) simulate 10,000 datasets coming from sensitive periods with different σ values ranging from 1 to 6, and add Gaussian noise to each simulated dataset based on the noise estimate in the control group, (2) fit the sensitive period model to each simulated dataset based on non-linear optimization, (3) calculate the Pearson correlation between age and MD change for each dataset and, (4) calculate the reliability of the fitted parameters and correlation coefficients. This simulation demonstrates that if the sensitive period for plasticity in the reading circuitry closes before 14 years of age, then we would have a high likelihood of detecting a significant correlation (>90%) and would be able to accurately estimate parameters of the sensitive period model. If the sensitive period extends through adolescence, then the upper bound on our parameter estimates would be unreliable. For example, if the true sensitive period is σ=5, meaning that the window for plasticity closes around 17 years of age, then we would still be able to infer that the sensitive period is open through elementary school (lower bound on confidence interval), but could not reliably infer the upper bound. In this case, we would also not have sufficient statistical power to detect a significant correlation (power = 0.7 at α= 0.05 for σ=5).

### Participants

The intervention group included 34 children (14 female), ranging in age from 7.2 to 12.8 years, who participated in an intensive summer reading program. Subjects were recruited based on parent report of reading difficulties and/or a clinical diagnosis of dyslexia. A total of 132 behavioral and MRI sessions were carried out with this group (more details on the participants can be found in our previous work (Huber et al., 2018)).

An additional 66 behavioral and MRI sessions were conducted in a non-intervention control group with 25 participants, who were matched for age but not reading level. These subjects were recruited as a control group to assess the stability of our measurements over the repeated sessions and this stability estimate was used as an estimate of noise in our simulations (**Figure 1**). Control subjects participated in the same experimental sessions but did not receive the reading intervention. Twelve of these control subjects had typical reading skills (5 female), defined as a score of 85 or greater on the Woodcock Johnson Basic Reading composite and the TOWRE Index. All subjects completed a battery of reading tests prior to the intervention period to confirm parent reports and establish a baseline for assessing growth in reading skill.

All participants were native English speakers with normal or corrected-to-normal vision and no history of neurological damage or psychiatric disorder. We obtained written consent from parents, and verbal assent from all child participants. All procedures, including recruitment, consent, and testing, followed the guidelines of the University of Washington Human Subjects Division and were reviewed and approved by the UW Institutional Review Board.

### Reading intervention

24 intervention subjects reported in Huber et al., 2018 were enrolled in 8 weeks of the *Seeing Stars: Symbol Imagery for Fluency, Orthography, Sight Words, and Spelling* (Bell, 2013) program at three different *Lindamood-Bell Learning Centers* in the Seattle area. An additional replication cohort of 10 subjects was run using the exact same training protocol at the University of Washington campus. The intervention program consists of directed, one-on-one training in phonological and orthographic processing skills, lasting four hours each day, five days a week. The curriculum uses an incremental approach, building from letters and syllables to words and connected texts, emphasizing phonological decoding skills as a foundation for spelling and comprehension. A hallmark of this intervention program is the intensity of the training protocol (4 hours a day, 5 days a week) and the personalized approach that comes with one-on-one instruction.

### Experimental sessions

Subjects participated in four experimental sessions separated by roughly 2.5-week intervals. For the intervention group, sessions were scheduled to occur before the intervention (baseline), after 2.5 weeks of intervention, after 5 weeks of intervention, and at the end of the 8-week intervention period. For the control group, sessions were scheduled to follow the same schedule while the subjects attended school as usual. This allowed us to control for changes that would occur due to typical development and learning during the school year. However, some of the parents of the subjects in the control group did not want their children to participate in all four scan sessions. In this case, the children were scheduled for two scan sessions and the timing of the scan sessions was determined such that, on average, we would have equal sampling of the different time intervals (∼3, ∼5 and 8 weeks).

In addition to MRI measurements, described in greater detail below, we administered a battery of behavioral tests in each experimental session. These included sub-tests from the Wechsler Abbreviated Scales of Intelligence (WASI), Comprehensive Test of Phonological Processing (CTOPP-2), Test of Word Reading Efficiency (TOWRE-2) and the Woodcock Johnson IV Tests of Achievement (WJ-IV).

### MRI acquisition and processing

All imaging data were acquired with a 3T Phillips Achieva scanner (Philips, Eindhoven, Netherlands) at the University of Washington Diagnostic Imaging Sciences Center (DISC) using a 32-channel head coil. An inflatable cap was used to minimize head motion, and participants were continuously monitored through a closed-circuit camera system. Prior to the first MRI session, all subjects completed a session in an MRI simulator, which helped them to practice holding still, with experimenter feedback. This practice session also allowed subjects to experience the noise and confinement of the scanner prior to the actual imaging sessions, to help them feel comfortable and relaxed during data collection.

Diffusion-weighted magnetic resonance imaging (dMRI) data were acquired with isotropic 2.0mm^3^ spatial resolution and full brain coverage. Each session consisted of 2 DWI scans, one with 32 non-collinear directions (b-value = 800 s/mm^2^), and a second with 64 non-collinear directions (b-value = 2,000 s/mm^2^). Each of the DWI scans also contained 4 volumes without diffusion weighting (b-value = 0). In addition, we collected one scan with 6 non-diffusion-weighted volumes and a reversed phase encoding direction (posterior-anterior, PA) to correct for EPI distortions due to inhomogeneities in the magnetic field. Distortion correction was performed using FSL’s *topup* tool (Andersson et al., 2003; Andersson and Sotiropoulos, 2016). The PA images were used to estimate the distortion field but were removed from all further analyses due to differences in mean signal intensity between the AP and PA images. Inclusion of the PA images in the kurtosis model fitting led to higher diffusivity estimates in our previous work (Huber et al., 2018), but all the results are equivalent whether or not these images are included.

Additional pre-processing was carried out using tools in FSL for motion and eddy current correction, and diffusion metrics were fit using the diffusion kurtosis model as implemented in DIPY (Garyfallidis et al., 2014). Data were manually checked for imaging artifacts and excessive dropped volumes. Given that subject motion can be especially problematic for the interpretation of group differences in DWI data (Koldewyn et al., 2014; Yendiki et al., 2014), data sets with mean slice-by-slice RMS displacement > 0.7mm were excluded from all further analyses. Datasets in which more than 10% of volumes were either dropped or contained visible artifact were also excluded from further analysis.

### White matter tract identification

Fiber tracts were identified for each subject using the Automated Fiber Quantification (AFQ) software package (Yeatman et al., 2012b), after initial generation of a whole-brain connectome using probabilistic tractography (MRtrix 3.0) (Jeurissen et al., 2011; Tournier et al., 2007). Fiber tracking was carried out on an aligned, distortion corrected, concatenated dataset including all four of the 64-direction (b-value = 2,000 s/mm^2^) datasets collected across sessions for each subject. This allowed us to ensure that estimates of diffusivity across session were mapped to the same anatomical location for each subject, since slight differences in diffusion properties over the course of intervention can influence the region of interest that is identified by the tractography algorithm.

### Quantifying white matter tissue properties

To detect intervention-driven changes in the white matter, we fit the diffusion kurtosis model as implemented in DIPY to the diffusion data collected in each session (Fieremans et al., 2011; Jensen and Helpern, 2010). The diffusion kurtosis model is an extension of the diffusion tensor model that accounts for the non-Gaussian behavior of water in heterogeneous tissue containing multiple barriers to diffusion (cell membranes, myelin sheaths, etc.). After model fitting, diffusion metrics were projected onto the segmented fiber tracts generated by AFQ. Selected tracts were sampled into 100 evenly spaced nodes, spanning termination points at the gray-white matter boundary, and then diffusion properties (mean, radial, and axial diffusivity (MD, RD, AD) and fractional anisotropy (FA)) were mapped onto each tract to create a “Tract Profile”. Here we focus on MD for three reasons: (1) MD is highly reliable and sensitive to changes in white matter tissue properties (De Santis et al., 2014); (2) MD was the primary metric of white matter plasticity reported in our previous work (Huber et al., 2018); (3) The simulations and power analysis in our pre-registration used the published MD data (https://github.com/yeatmanlab/Huber_2018_NatCommun).

## Results

### The magnitude of white matter plasticity does not vary with age

In our previous work we observed intervention-driven changes in white matter tissue properties that were distributed across a network of highly correlated white matter tracts (Huber et al., 2018, Figure 6). To summarize this spatially distributed plasticity, and to limit the number of statistical comparisons, we first conducted a principal component analysis of mean diffusivity (MD) values sampled at 100 nodes along the 18 major white matter fascicles. Values from all 18 tracts were concatenated in a matrix (34 subjects X 4 time points by 18 tracts X 100 nodes); columns were centered but not scaled. The first 5 components accounted for 55 percent of the variance in the data (PC1 = 27%, PC2 = 10%, PC3 = 7%, PC4 = 6%, PC5 = 5%). Since each component is a linear combination of MD values from different white matter tracts, analyzing each subject’s change in component scores is akin to averaging MD values across a collection of tracts with similar diffusion properties (see **Figure S1** for a visualization of PC1). Using a linear mixed effects LME model to analyze changes in MD as a function of intervention hours (fixed effect of intervention time in hours and random effect of subject), we find significant intervention driven change in PC1 (t(112) = −3.10, p = 0.0024) and PC4 (t(112) = 3.00, p = 0.0034). However, the amount of change observed in an individual subject is not dependent on that subject’s age. Specifically: (1) fitting an LME model with age, time (intervention hours), and their interaction, indicated that the interaction was not significant for any of the five PCs (all p-values > 0.16); (2) the correlation between age and individual growth rates (change in MD per hour of intervention estimated as a linear fit to each individual’s data) were near zero (PC1 r = 0.03, p = 0.90; PC2 r = −0.08, p = 0.69; PC3 r = −0.28, p = 0.15; PC4 r=0.05, p = 0.81; PC5 r = −0.21, p = 0.27); (3) a group comparison for average growth rates in younger (<9 years of age) versus older subjects showed no group differences (all p-values > 0.20). Computing the Bayes factor (BF, based on (Wetzels and Wagenmakers, 2012)) provides moderate support for the null hypothesis that the magnitude of intervention driven white matter plasticity does not depend on age (PC1 BF = 0.15; PC2 BF = 0.16; PC3 BF = 0.40; PC4 BF = 0.15; PC5 BF = 0.26).

### Post-hoc node-based analysis of age dependent plasticity

Given that spatially distributed plasticity (in terms of principal components) observed over the intensive reading intervention is equivalent for older versus younger subjects, we next undertook a post-hoc analysis (not defined in the pre-registration) to search for regions in the white matter that show a relationship between age and intervention driven growth rate. In this analysis we compute MD growth rates for each of the 100 nodes along each of the 18 white matter tracts and correlate these growth rates with subject age. Figure 2 shows a rendering of the 18 white matter tracts colored based on the average intervention-driven change in MD at each node (**Figure 2A**), and the relationship between age and MD change at each node (**Figure 2B,C**).

**Figure 2:**
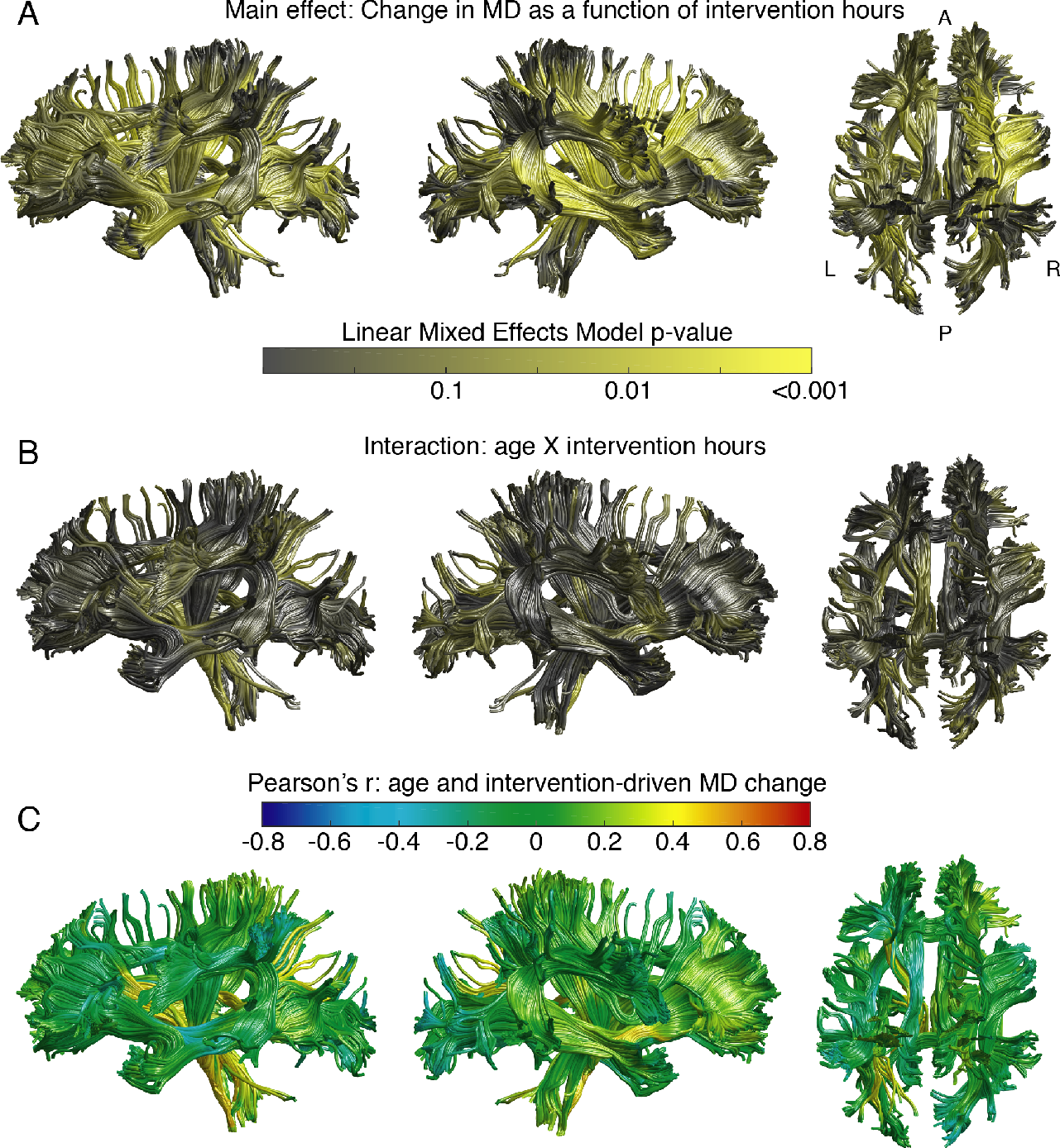
Intervention-driven white matter plasticity does not depend on age. A Linear mixed effects model predicting changes in mean diffusivity (MD) as a function of intervention hours, age and their interaction was fit at each node along each white matter tract. (A) Highly significant intervention-driven changes in MD were distributed throughout the white matter (main effect of intervention hours). (B) The age by intervention hours interaction was not significant throughout most of the white matter. Focal regions of the corticospinal tract were significant without a correction for multiple comparisons, but the extensive network seen in panel A largely showed no relationship between age and plasticity. Histograms of p-values for each tract are shown in Figure S3. (C) Correlation coefficients (Pearson’s r) relating each subject’s age to their rate of intervention-driven MD change are shown as an index of effect size. Correlations are mostly distributed around zero.

This exploratory analysis revealed effects that were isolated to small regions on a few tracts (e.g., ventral portion of the left corticospinal tract, minimum p = 0.006 uncorrected, **Figure 2B,C,** see also **Figure S3**). However, like a voxel-based analysis (VBA), asserting statistical significance based on an exploratory node-based analysis requires a correction for multiple comparisons (accounting for the correlation structure in the data as nodes are not independent). No regions showed a strong relationship between plasticity and age, and these effects would not survive a multiple comparison correction (see **Figure S2** and **Figure S3** for histograms of p-values). Thus, while we cannot rule out the possibility that there exists a specific focal region that loses its capacity for plasticity over the course of elementary school, the extensive networks of regions showing intervention-driven change (**Figure 2A**) do not show interactions with subject age (**Figure 2B**).

### The time-course of white matter plasticity does not vary with age

Given that the magnitude of white matter plasticity does not change as a function of age, we next turn to our second hypothesis that younger subjects will show more rapid changes compared to older subjects, irrespective of the absolute magnitude of change. We once again begin by focusing on the first five principal components and compute the intervention driven growth rate in MD (change/hr) observed between the first two measurements sessions (which were separated by, on average, 26 days). We find that the rate of change in MD during the beginning of the intervention does not depend on age (spearman correlation, PC 1 r = 0.03, p = 0.89; PC 2 r = −0.16, p = 0.41; PC 3 r = 0.06, p=0.77; PC 4 r = −0.21, p = 0.29; PC5 r = 0.11, p = 0.58). Finally, we calculate the percentage of overall change that occurs during the first three weeks of intervention by using linear regression to interpolate each subject’s MD value at 60 hours of intervention, subtracting the subject’s baseline MD value, and dividing this difference by the difference between baseline and post-intervention MD. The point of this interpolation is to control for the fact that there was some variability among subjects in the timing of measurement session 2. Overall, we once again find that the percentage of overall change occurring within the first three weeks of the intervention does not depend on age: there was a marginally significant correlation for PC2 (r = −0.43, p = 0.03 uncorrected), however, PC2 did not show a significant main effect of the intervention. Visualizing the time-course of change in older versus younger subjects reveals very similar growth trajectories.

### Older children and younger children show equivalent improvements in reading scores

Finally, to ascertain if the efficacy of the intervention (in terms of behavioral improvements) changes as a function of age, we perform a similar set of analyses on the behavioral measures. Previous intervention studies have demonstrated greater gains in younger, compared to older children (Lovett et al., 2017), though no study has examined age effects in an intensive (4 hour a day), one-on-one intervention program such as the one employed here. This is considered a post-hoc analysis because it was not specified in our pre-registration. **Figure 3b** shows intervention-driven growth trajectories for the Woodcock Johnson Basic Reading Skills composite score. Qualitatively, the growth trajectories are very similar for each group. Indeed, there was not an age by time interaction for any of the reading measures. For the Woodcock Johnson Basic Reading Skills composite there was a highly significant main effect of the intervention (t(110) = 8.73, p = 3.41 × 10^−14^), a significant main effect of age (older subjects in our sample were, on average, more severely impaired readers than the younger subjects: t(110) = −3.43, p = 8.4 × 10^−4^), but no interaction (t(110) = −0.76, p = 0.44). For the Test of Word Reading Efficiency composite score, there was a highly significant main effect of the intervention (t(110) = 6.23, p = 7.3 × 10^−9^), a small but significant main effect of age (t(110) = −2.31, p = 0.02), but no interaction (t(110) = 0.01, p = 0.99).

**Figure 3:**
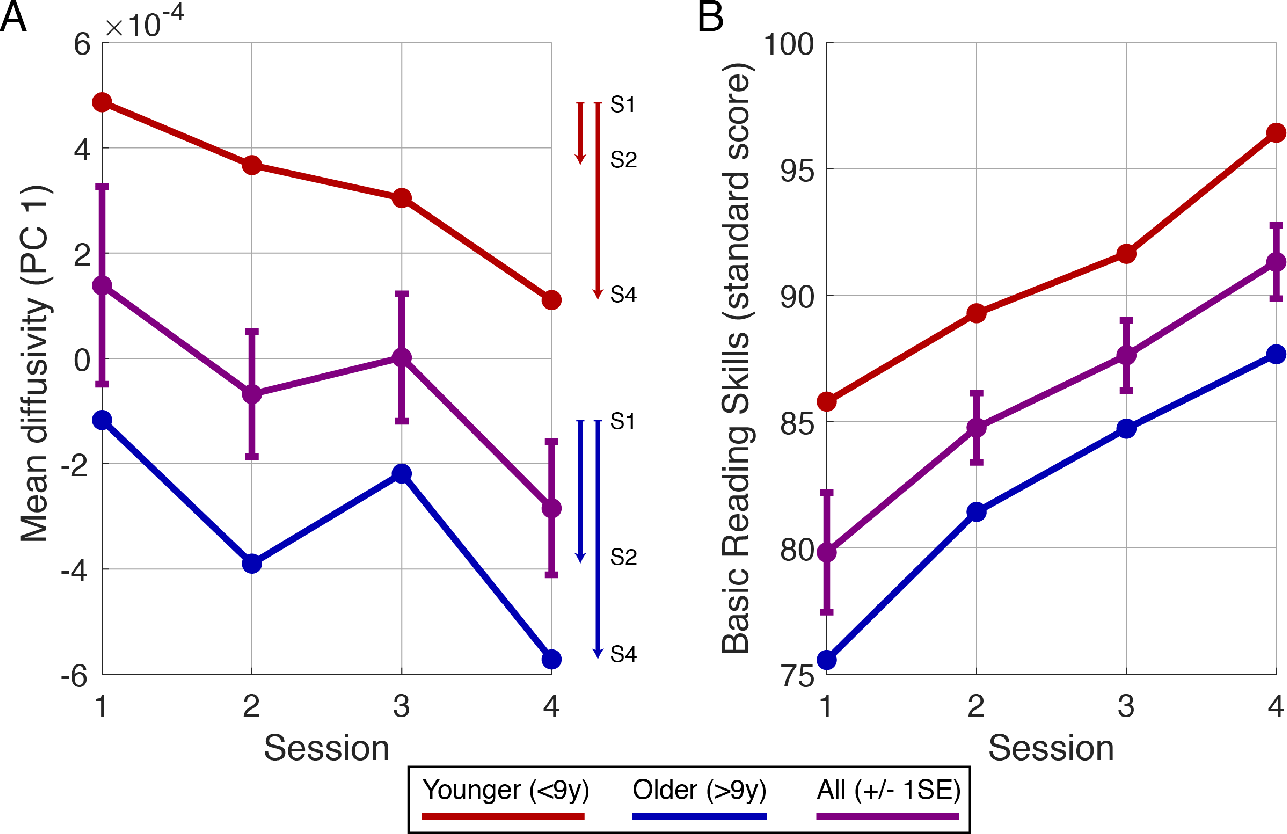
Time-course of white matter plasticity and reading skill growth. (A) The time-course of change in mean diffusivity (MD) for the first white matter principal component (PC1) is plotted for the whole sample (purple), younger subjects (red) and older subjects (blue). Initial MD is lower in the older subjects, compared to the younger subjects, reflecting known maturational effects. But the time-course of plasticity is equivalent for the two groups. The vectors on the right side of the plot show the magnitude of change between the first two sessions and after completion of the intervention. Contrary to our hypothesis, the younger subjects showed slightly less change between the first two sessions and slightly less change overall (no significant differences). (B) In terms of behavioral improvements, the time-course and magnitude of reading gains were equivalent for both groups.

## Discussion

We combined an intensive, one-on-one reading intervention program with longitudinal diffusion MRI measures of the white matter in a sample of children spanning early elementary school (7 years of age) through middle school (13 years of age) to determine whether a child’s propensity for white matter plasticity changes over these years of development. We found that, for the intervention program employed here, neither white matter plasticity nor reading growth rates diminish between first grade and middle school. These findings stand in stark contrast to our predictions: (1) in terms of white matter plasticity, younger and older children show rapid and widespread changes with an equivalent magnitude and time-course; (2) in terms of improvements in reading skills, younger and older children show equivalent gains in reading accuracy and reading rate.

That the capacity for intervention-driven white matter plasticity does not appear to change over the course of elementary school comes as a surprise. The broader literature on reading interventions makes it clear that, on average, intervention programs are more effective in younger compared to older children (Lovett et al., 2017; Torgesen, 1998). Thus, there was reason to believe that intervention-driven plasticity might diminish with age. Moreover, data from rodents has indicated a sensitive period for association cortices that closes at the onset of puberty, leading to the prediction of diminished plasticity over this age range for the circuitry supporting human high-level cognitive functions (Piekarski et al., 2017b). However, recent data on language learning in a large sample of 2/3 million English speakers revealed that the sensitive period for syntax learning extends later than expected; learning rates remain constant until 17 years of age at which point learning rates sharply decline (Hartshorne et al., 2018).

What is unique about the intervention employed here is the intensity (4 hours a day, five days a week), and the one-on-one delivery of the curriculum, which allows a skilled instructor to tailor the pace of the program to the individual student’s needs. These two factors might be critical for promoting both white matter plasticity and learning over a broad age range. In line with this idea, we see large behavioral effects across the full age range of our sample, which deviates from the expectation that intervention efficacy declines sharply with age. Determining the specific aspects of an intervention and, more broadly, the environmental factors that prompt large-scale changes in white matter tissue properties, is an important direction for future research.

The present findings are also subject to other interpretations. For example, in implementing a model of sensitive periods, we used a Gaussian function bounded at zero to be consistent with well-characterized critical periods that are present early in maturation (Hensch, 2005; Werker and Hensch, 2014). However, it is known from animal models that white matter plasticity is still present in the adult brain (Fields, 2015; Gibson et al., 2014; McKenzie et al., 2014; Mount and Monje, 2017). Moreover, human language learning is known to prompt white matter changes in the adult brain (Hofstetter et al., 2017; Schlegel et al., 2012). While we can be reasonably certain that there is not a dramatic change in white matter plasticity (as indexed by diffusion) across the age range sampled here, we cannot rule out the possibility of gradual declines beyond the sampled age-range. Nor can we rule out the possibility that a sensitive period occurs very early in elementary school (e.g., 5-7 years of age), with continued but diminished plasticity throughout childhood and into adulthood. Resolving a sensitive period that occurs early in elementary school would require a larger sample of kindergarten and first grade children, and testing the prediction of a sensitive period model with more parameters (e.g., adding a constant term to the Gaussian model) would require a larger sample across a broad age range. Furthermore, we focused on mean diffusivity due to its excellent sensitivity to changes in white matter structure (De Santis et al., 2014; Huber et al., 2018). However, mean diffusivity is also not biologically specific, and is affected by properties of axons, myelin, glia and vasculature; it is possible that specific cell types have specific sensitive periods that could be detected with a more biologically specific quantitative MRI measurement (Berman et al., 2017; Mezer et al., 2013; Stikov et al., 2011; Weiskopf et al., 2015; Yeatman et al., 2014).

What are the implications of these findings for literacy education? First, these data prompt a new interpretation of white matter differences reported in children with dyslexia (Ben-Shachar et al., 2007; Ozernov-Palchik and Gaab, 2016; Vandermosten et al., 2012; Wandell and Yeatman, 2013). When children are provided a high-quality educational experience, the white matter is much more plastic than often presumed; therefore, observed differences in white matter organization are just as likely to be a consequence, as oppose to a cause, of differences in literacy achievement. Second, intervention programs can still be highly effective in older children, even ones with extreme reading difficulties. However, early intervention should still remain the gold standard: not intervening to remediate reading difficulties has well-known consequences in terms of increased risk for anxiety and depression (Nelson and Harwood, 2011), and poor academic outcomes (struggles with reading tend to exert a negative impact across many academic domains; Snow et al., 1998; Snowling, 2013; Torgesen, 2004, 1998). Finally, this study did not measure long-term outcomes and it is possible that short-term intervention is more effective at eliciting long term change (in white matter and behavior) in younger brains.

In sum, this work highlights the powerful influence that education exerts on white matter development throughout elementary school. We did not discover a sensitive period for white matter plasticity. With the growing literature on neuroanatomical correlates of learning disabilities, our findings underscore the importance of investigating the conditions under which different neural systems are plastic or stable.

## Acknowledgements

We would like to thank Rebecca Saxe, Hilary Richardson and Timothy Verstynen, Shai Berman and Aviv Mezer for comments and discussion of the preprint. We would also like to thank Patrick Donnelley and Emily Kubota for their role in data collection. This work was supported by NSF/BSF BCS #1551330, NICHD R21HD092771, NICHD 2P50HD052120-11, and Jacobs Foundation Early Career Research Fellowship to J.D.Y.

## Supplementary Figures

**Figure S1:**
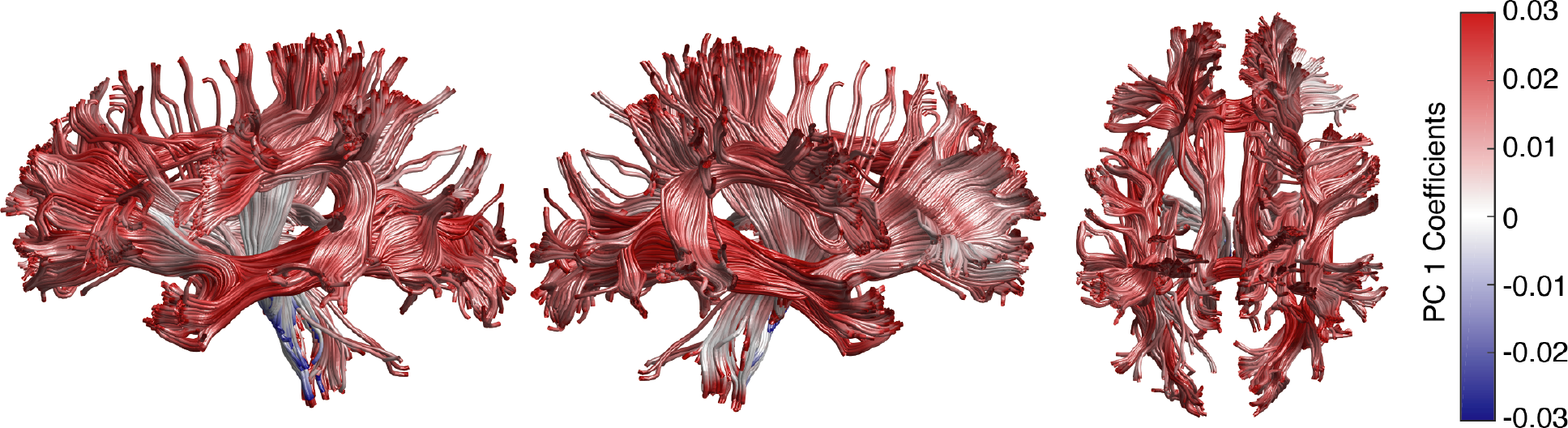
Visualization of the first principal component of mean diffusivity values. Principal component analysis (PCA) was used for dimensionality reduction. Each component is a weighted sum of mean diffusivity values sampled across the white matter. Thus, change in a subject’s component score over the intervention indexes change across this widespread network of correlated diffusivity values. Component loadings for the first principal component are visualized as a colormap.

**Figure S2:**
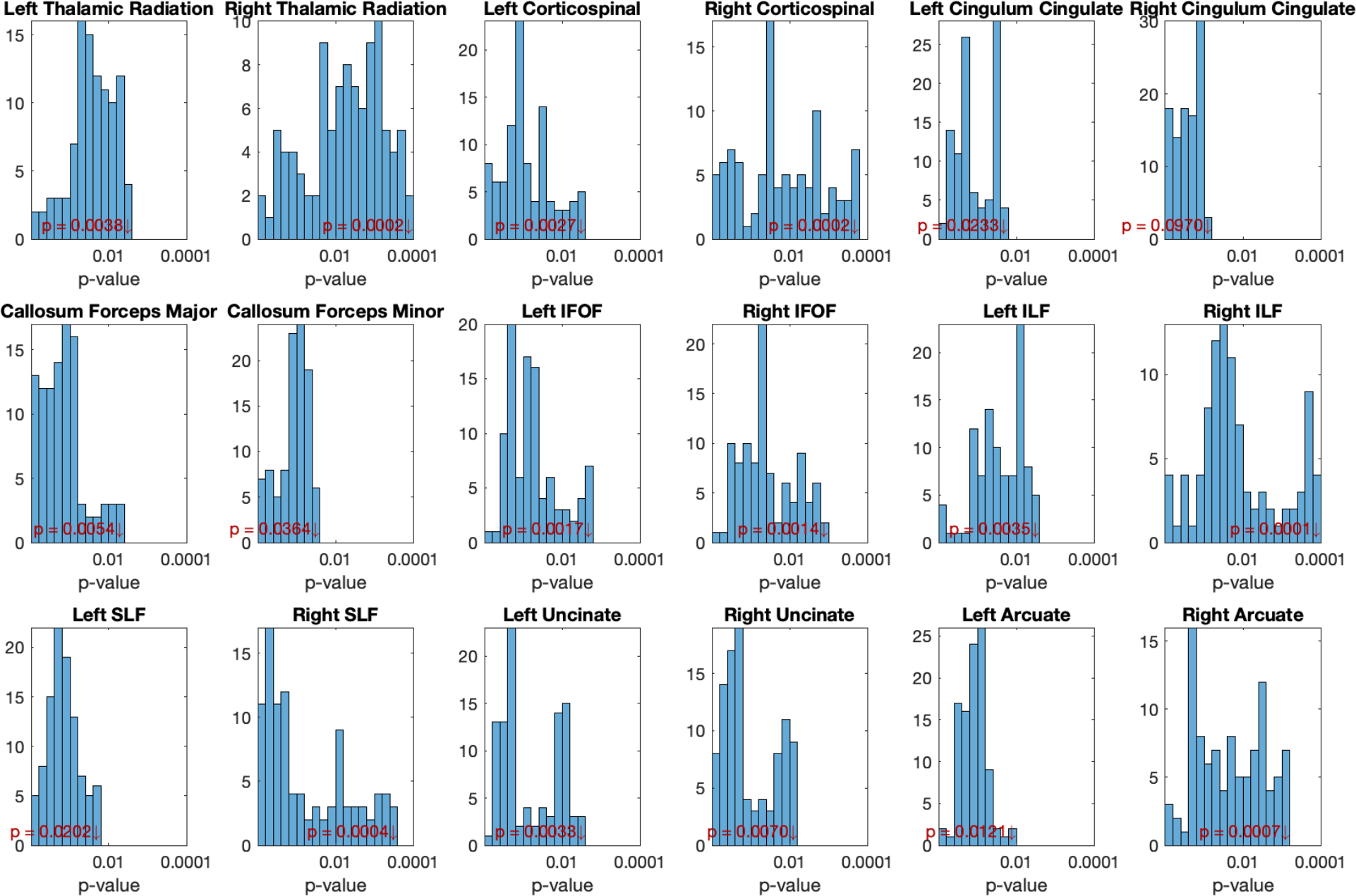
Main effect of intervention time. Histograms of p-values for each tract rendered in Figure 2A. P-values are given for the main effect of time in the linear mixed effects model of mean diffusivity values as a function of intervention time. Red arrow denotes the smallest p-value for each tract.

**Figure S3:**
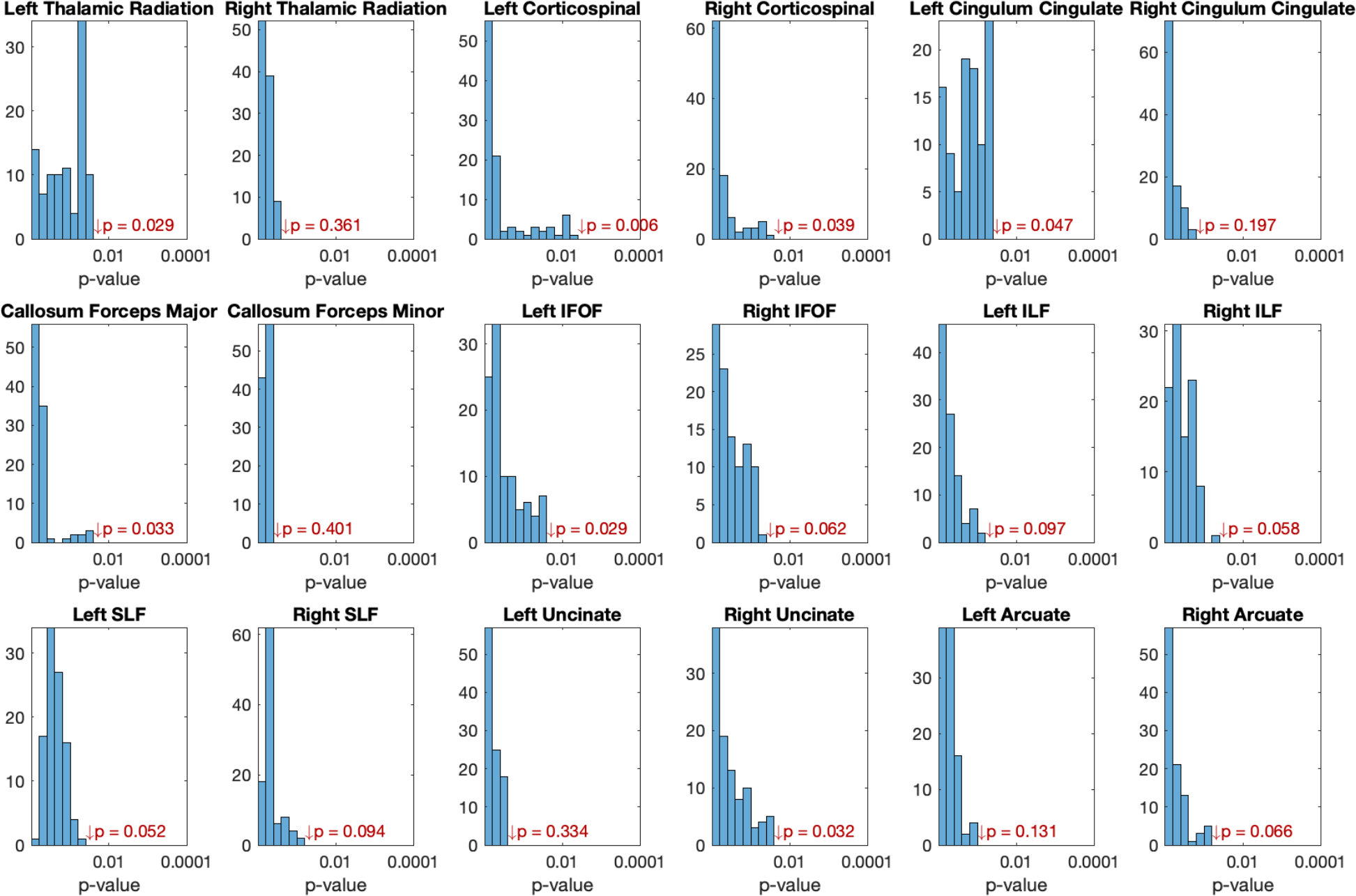
Age by Time interaction. Histograms of p-values for each tract rendered in Figure 2B. P-values are given for the interaction term in the linear mixed effects model: change in plasticity as a function of age. Some tracts show regions that surpass a threshold of p<0.05 but would not survive a correction for multiple comparisons. Red arrow denotes the smallest p-value for each tract.

